# Spatio-temporal patterns of ovarian development and *VgR* gene silencing reduced fecundity of parthenogenetic *Artemia*

**DOI:** 10.1101/2023.05.03.539339

**Authors:** Hu Duan, Xuanxuan Shao, Wei Liu, Jianhai Xiang, Namin Pan, Xuehui Wang, Guoru Du, Ying Li, Jiaping Zhou, Liying Sui

## Abstract

Halophilic zooplankton brine shrimp *Artemia* has been used as an experimental animal in multidisciplinary research. However, the reproductive patterns and regulation mechanisms of *Artemia* remain a mystery. In this study, the ovarian development process of parthenogenetic *Artemia* (P. *Artemia*) was distinguished into five stages, and oogenesis or egg formation process was identified with six phases. The mode of oogenesis was assumed polytrophic. Moreover, we traced the dynamic translocation of germline stem cells (GSCs) by EdU labeling, elucidated several key cytological events in oogenesis through HE staining and florescent image. Notably, distinguished from the ovary structure of insects and crustaceans, P. *Artemia* germarium originated from ovariole buds and located in the base of the ovarioles. RNA-seq based on five stages of ovarian development revealed that 2657 up-regulated genes related to reproduction were obtained by pair-to-pair comparison of RNA-seq. Furthermore, *Gbb, Dpp, piwi, vasa, nanos, VgA* and *VgR* associated with GSCs recognition and reproductive development genes were screened and also verified by qPCR. Silencing of the *VgR* gene in P. *Artemia* (*PA-VgR*) at stage L of ovarian development led to a low level of gene expression (<10%) within five days, which resulted variations of oogenesis-related gene expression and significantly inhibited vitellogenesis, impeded oocyte maturation, and eventually decreased the offspring number. In conclusion, we illustrated the patterns of ovarian development, outlined the key spatio-temporal features of oogenesis and established a platform for gene function research using P. *Artemia* as an experimental animal. These results lay the foundation for studying the reproductive biology of invertebrates, may also provide a novel insight in the reproductive regulation mechanism of aquatic animals in extreme environments.

**SUMMARY STATEMENT:** Artemia is a novel experimental platform and potential “model” crustacean for studying specific process and mechanism of reproduction.

## INTRODUCTION

Reproductive development is vital in body developmental processes and propagation of multicellular organisms (Roy et al., 2018). The sequentially coordinated events of the ovarian development and oogenesis (also known as egg formation) are critical for female reproduction (Ruan et al., 2020). Egg formation in oviparous females, especially in classical “model” organisms and some insects have been well studied (Shapiro-Kulnane et al., 2015; Leyria et al., 2021), but the subject of evolutionary developmental biology (evo-devo) has become an emerging field that requires a broader picture across tremendous biodiversity (Ponz-Segrelles et al., 2018; Zhao et al., 2021). In general, ovarian development in bisexual species includes the process of mature follicle generation, estrogen secretion, egg production, fertilization and other important processes (Lu et al., 2018), which accompanies by the occurrence of successive cytological events. In *drosophila*, oogenesis requires extensive interaction between somatic and germline stem cells, and these developmental events are accomplished in the ovaries (Fu et al., 2015b). Oogenesis mainly includes primordial germ cell (PGC) formation, oocyte development, differentiation and maturation, which were controlled and regulated by relevant genes and signaling pathways (Xie et al., 2017). As a hot subject in developmental biology and reproductive biology, oogenesis has been extensively studied in mammals, insects and a few crustaceans (Neaves and Baumann, 2011; Herawati et al., 2020; Zhang et al., 2022). However, the lack of knowledge on the pattern of reproductive processes and molecular mechanisms in invertebrates, especially for the species living in extreme habitats (*e*.*g*. hypersaline, hypoxia, etc.), restrains the comprehensive knowledge and understanding of reproductive evolution.

Hypersaline environment provides extraordinary habitat for all domains of life on Earth since the Precambrian period (Gajardo and Beardmore, 2012). Brine shrimp *Artemia* affiliate to Crustacea; Pancrustacea; Branchiopoda, thriving in hypersaline water body with salinity up to 20%. *Artemia* has several biological advantages and special life phenomena, such as transparent body, asexual and bisexual reproduction modes, oviparity and ovoviviparity breeding strategy as well diapause and de-diapause life cycles (MacRae, 2016). And thus, *Artemia* have been used as an experimental animal in environmental toxicology, oncology, epigenetics and other research fields (Ye et al., 2017; Suman et al., 2020; Balamurugan and Balasubramani, 2022). Compared with bisexual *Artemia*, parthenogenetic *Artemia* (P. *Artemia*) impose on the special reproduction mode. In other words, the female accomplishes a process of sexual reproduction in which the unfertilized egg develops without the contribution of the male gamete, which ensures to produce offspring that develops from unfertilized embryos are considered “full clones” with the same genotype and are genetically identical to their mother, as well repeatability of the experiments. In this regard, P. *Artemia* can be considered as a potential animal model for investigating the organizational basis and reproductive molecular mechanism of animals adapting to the harsh environment (Zhao et al., 2021).

RNA-Seq approach is widely used to analyze the gene networks associating with development processes at the transcriptional level and to identify the development relating genes in a variety of invertebrates (Qiao et al., 2017; Wang et al., 2019). By analyzing the interaction between metabolites and transcriptomes, the reproductive relating genes that regulate cell division and hormone signal transduction in the course of ovarian development were screened in fresh water prawn *Macrobrachium nipponense* (Zhang et al., 2020). For oviparous animals, vitellogenesis is a prerequisite for organism oviposition and embryo development (Mitchell et al., 2019). Vitellogenin (Vg) is synthesized in hepatopancreas of crustaceans (Jia et al., 2013) and the fat body of insects (Zou et al., 2020). Vg is transported through the blood, binds to vitellogenin receptors (VgR) located in the plasma membrane of oocyte to form Vg/VgR complex, and eventually enters oocyte through endocytosis (Chen et al., 2020). Moreover, findings that *VgR* knockout or mutation can inhibit the oocytes maturation of insects or cause sterility of birds, also indicating that VgR plays a crucial role in ovarian development of oviparous animals (Lin et al., 2013; Ali et al., 2017). In addition, the researchers have also focused on other genes that play a key role in oogenesis, such as *Dpp* and *Gbb* in BMP signaling pathways, as well as *vasa, nanos* and *piwi* in reproductive stem cells. Among them, *Dpp* and *Gbb* mainly promote the growth of follicular cells or regulate the synthesis of juvenile hormone (JH) by inhibiting the gene encoding JH acid o-methyltransferase (Daans et al., 2008; Ishimaru et al., 2016). Silencing of *vasa*, a marker of reproductive stem cells, changed the morphology of both ovaries and oocytes, and reduced the number of eggs in Japanese bloodsucking worm *Schistosoma japonicum* (He et al., 2018). Moreover, *nanos* and *piwi* are important in regulating the expression of germline stem cells (Cox et al., 1998). So far, no study has been conducted on the regulatory genes, pathways, functions and molecular mechanisms of genes in process of P. *Artemia* oogenesis.

Our study reported the morphological and tissue characteristics in ovarian development process, clarified the cytological details of oogenesis, and screened some candidate genes that might involve in ovarian development and oogenesis in P. *Artemia* with focusing on the function of *PA-VgR* gene in process of oogenesis. Our goal in this study is to provide novel insights for understanding the mechanism of ovarian development in extreme environmental reproductive biology, but also establish a platform for the study on a special life process of aquatic invertebrates.

## MATERIAL AND METHODS

### Artemia rearing

P. *Artemia* cysts from Aibi lake, China was obtained from Asian Regional Artemia Reference Center (ARARC), Tianjin University of Science and Technology, China. About 0.2 g cysts were incubated in a conical tube containing 200 mL dilute brine water (salinity of 3%) under constant hatching condition (28□, 2000 Lux illumination and continuous aeration). After 24 h, *Artemia* nauplii were collected and transferred to a rectangular plastic tank containing 16 L diluted brine (salinity of 7%). The animals were fed twice a day with microalgae *Chlorella* sp. Water was renewed once a week. The ovaries morphology of P. *Artemia* were observed under a stereoscope (SZ680, China) when the oocyst primordia appeared.

### Chemical dyeing

To examine the structural characteristics in process of the ovarian development, P. *Artemia* were fixed in 4% paraformaldehyde (Acros Organics, Lot: A0372619) diluted with PBS (6.6 mM Na_2_HPO_4_/KH_2_PO_4_, 150 mM NaCl, pH 7.4) for 2 h. The ovaries were dissected using ophthalmic forceps (Spi-Swiss) and incubated in 0.5% Triton X-100 (Sigma, Lot: SLBP6453V) for 25 min. After PBS rinsing, the ovaries were stained with Phalloidin (Solarbio, 1029O011) and Hoechst 33342 (ThermoFisher, Lot: 1681305) for 4 h. Anti-fluorescence quenching agent (Beyotime, P0126) was added after PBS rinsing to avoid fluorescence quenching. And the tissue samples were observed under a fluorescence microscope (Nikon Ti-E, Japan).

### HE staining

To characterize the histological and cytological features in process of the ovarian development, the P. *Artemia* individuals at five distinguished developmental stages were collected, and immediately fixed in 4% paraformaldehyde at 4□ for 24-48 h (Zou et al., 2020). Briefly, the tissues were dehydrated, permeated, and immersed in xylene and paraffin wax (1:1, v:v) (Bellancom, Lot: DGBF4846V) over night, and then were dewaxed and stained with hematoxylin-eosin (BKMAM, Lot:20220520; 20220428). And the tissue samples were observed under microscope (Nikon Ti-E, Japan).

### Immunofluorescence antibody labeling

To describe the dynamic migration pathway of germline stem cells during oogenesis, the P. *Artemia* ovaries were dissected as clean as possible using forceps, and incubated with a mixture of primary antibodies in the cell incubator (Corning) for 2 h. The mixture composed of 2× EdU working solution (ThermoFisher, Lot: 2261449), DMEM medium (Gibco, Lot: 1897090), penicillin, streptomycin and neomycin (Gibco, Lot: 1894168). The tissues were fixed in 4% paraformaldehyde for another 2 h. After rinsing with PBS, the tissues were sealed with 3% BSA (ThermoFisher, Lot: SF248474) for 20 min. And 0.5% Triton was used to increase the permeability of antibodies to the cell membrane. Secondary antibodies which conjugated with Alexa Fluor 488 (ThermoFisher, Lot: 2261449) were incubated (1:500, v:v) with the tissues for 2 h in dark and moist condition. Lastly, Hoechst 33342 were incubated with samples at room temperature for 30 min in a ratio of 1:2000 (v:v). Anti-fluorescence quench agent was added after cleaning. Images were obtained by a laser confocal microscope (Zeiss 980, Germany).

### RNA extraction and Illumina sequencing and data analysis

The P. *Artemia* ovaries at five developmental stages were dissected in cold liquid nitrogen. Thirty individuals at the same developmental stage were pooled. RNA extraction was performed with RNAiso Plus reagent (TaKaRa, AKF0727A). Transcriptome library construction and subsequent sequencing analysis were completed by Novogene Technology Co., Ltd, China.

The original data obtained by high-throughput sequencing were filtered. The functional unigenes were annotated by seven databases including GO (http://www.geneontology.org/) and KEGG (https://www.kegg.jp/), etc. The gene structural domains were analyzed using SMART program. The phylogenetic tree was constructed by a maximum likelihood method using MEGA7 with bootstrap of 1000 replicates.

### RT-qPCR analysis

To explore the spatio-temporal expression patterns of reproductive genes in P. *Artemia*, total RNA was extracted from the target tissues (intestine, cerebral ganglion and ovary) at five developmental stages using RNAiso Plus reagent (TaKaRa, Lot: AKF0727A). cDNAs were synthesized from 1μg of total RNA (ThermoFisher, Lot: A48571). RT-qPCR was performed using a TB Green®Premix Ex Taq™II (TaKaRa, Lot: AL51019A), and β-actin gene was used as a reference gene. The primers were designed by Primer 5 (Table 1). Each reaction contained three technical replicates as well as template free control.

**Table 1.**
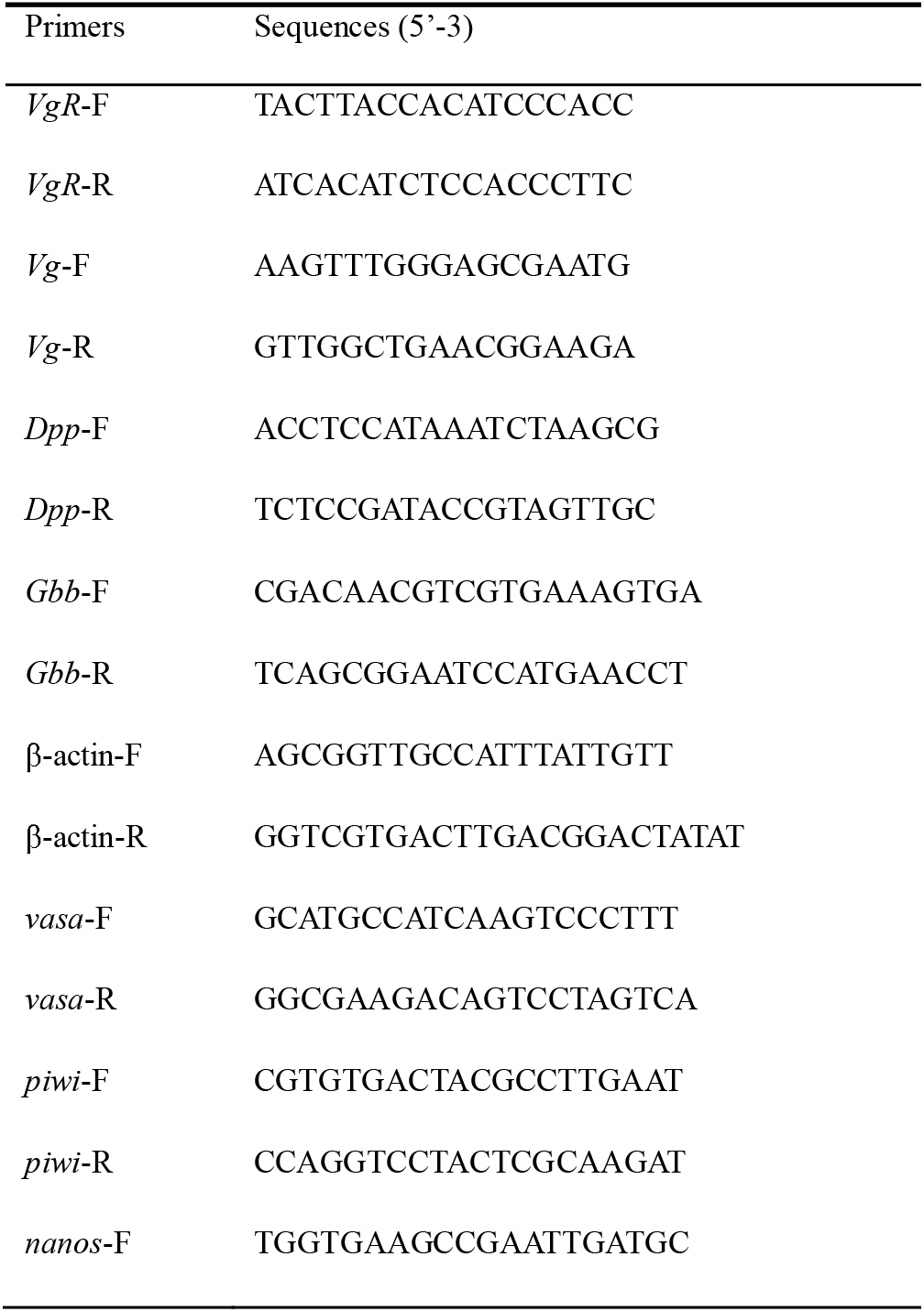
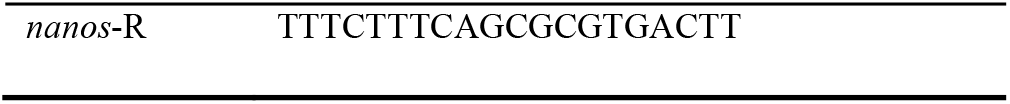
Primers used in the functional gene study.

### RNAi and phenotypic characterization

Microinjection needles were prepared using micropipette puller (Narishige PC-10, Japan) and 1.0 mm capillary tubes (VitalSense Scientific Instruments Co., Ltd. Lot: B100114N). Double strand RNA (dsRNA) of *VgR* and *EGFP* were synthesized using T7 RiboMAX™ Express RNAi System (Promega, USA). Primers for dsRNA synthesis were summarized in Table 2. After anesthetized on ice, about 2000 ng dsRNA was injected into the ventral side of the second abdominal segment of P. *Artemia* using a microinjector (Nikon Eclipse Ti, Japan) fitted with an injection needle. P. *Artemia* in the control group were treated with the same dose and volume of ds*EGFP*. The ovaries were sampled on day 1^st^, day 3^rd^ and day 5^th^ after dsRNA injection, and total RNA was extracted. The knockdown efficiency and gene variation were determined by qRT-PCR.

**Table 2.**
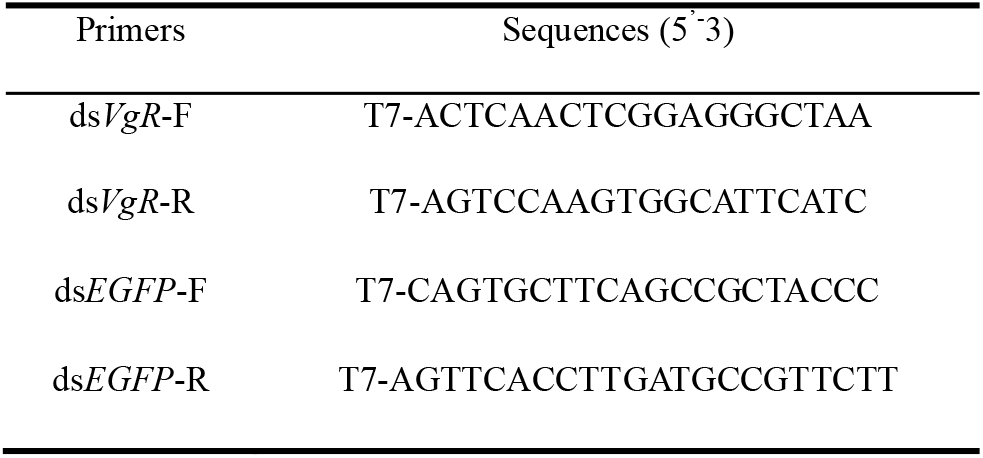
Primers for dsRNA synthesis.

For phenotypic characterization, the ovaries were collected when more than 80% P. *Artemia* had reached the objective developmental stages, and the ovary morphologies and cytological features were determined as described previously in session 2.3. The morphology and cytologic variation of the ovaries were observed under an inverted fluorescence microscope (Nikon Ti-E, Japan).

### Statistical analyses

Data were presented as mean ± standard deviation. Statistical significance was determined using one-way ANOVA followed by Duncan multiple comparisons at P<0.05 and P<0.01, respectively (SPSS26.0). The statistically significant difference between two groups was inferred using T-test.

## RESULTS

### Tissue characteristics in process of ovarian development

After about 24 h hatching, most of the gastrula embryo of P. *Artemia* developed into Instar I nauplius stage which was considered as a starting point for characterizing the ovarian development progress (Fig.1, 0 dph, day post hatching). Within 7-10 dph, about 63% individuals gradually developed into the ovarian developmental period (stage □) with appearance of obvious pair of ovariole buds (dark arrows, *in vitro*) near to the anterior ventral surface of the body. At 15-17 dph, the ovaries developed to mature period, of which approximate 51% individuals were further distinguished into pro-mature stage (□), meta-mature stage (Ш) and ana-mature stage (□), according to the ovarian morphology and characteristics (i.e. ovariole shape, location and features of oocytes) (Fig.1, stage □-□). At 25-29 dph, the ovaries developed into ovulatory period (stage □), and the egg resided in the oocyst with no longer transparent as a result of yolk granule accumulation.

**Fig. 1.**
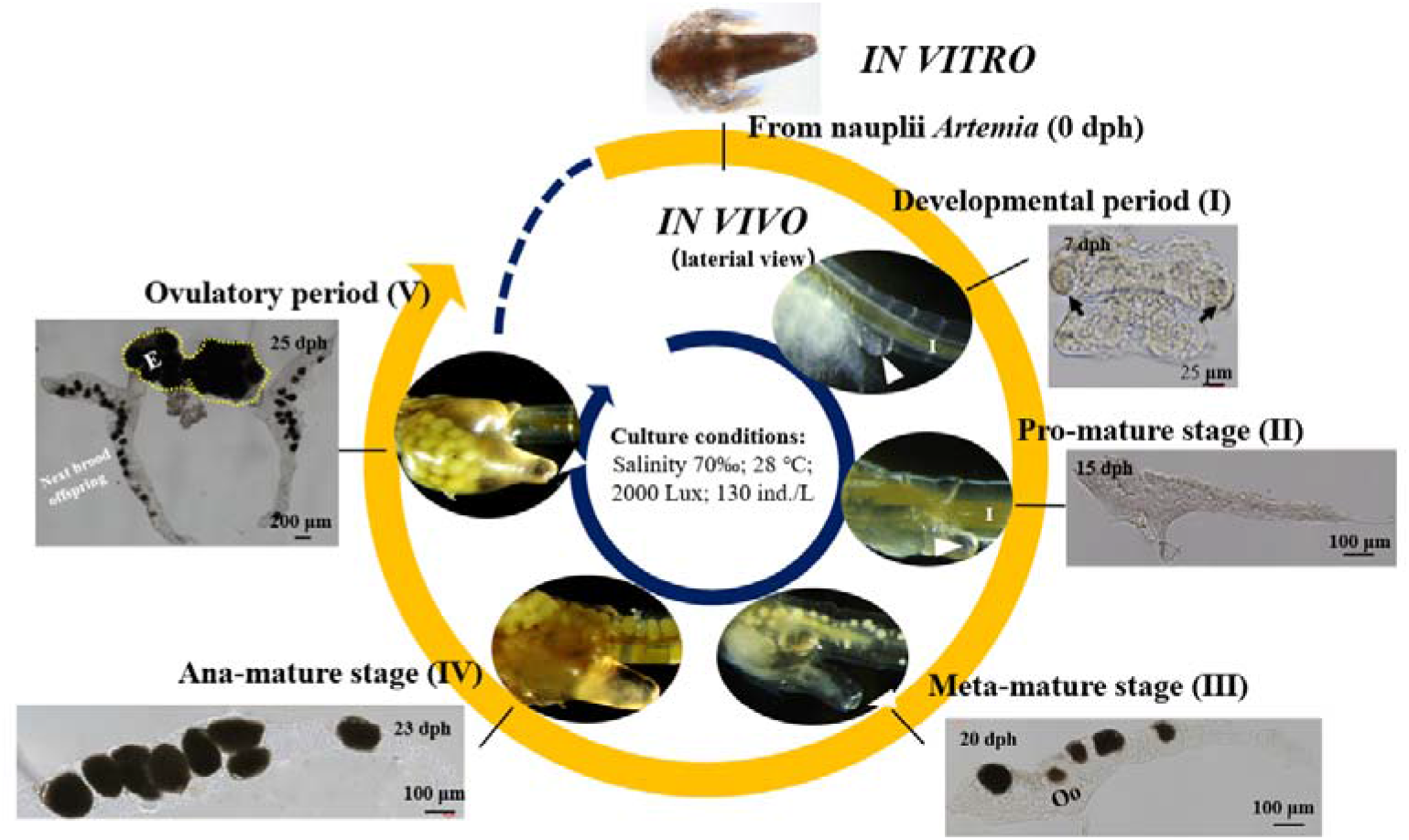
The temporal patterns and morphological characteristics in process of ovarian development of P. *Artemia*. dph: day post hatching; E: eggs; I: Intestine; Oo: oocyte; dark arrow: ovariole bud; triangular white arrow: ovulatory hole; yellow dot line: oocyst.

### Key cytological events of oogenesis

The cytological characteristics of the ovaries were revealed by Hoechst labelling and HE staining. The ovaries showed general bilateral symmetry through the entire ovarian developmental process (Fig. 2, I-1,2, white dot line). Based on the features of vitellogenesis, and accumulation and distribution of yolk granules observed under optical and fluorescence microscope, six phases of oogenesis (from oogonia to multicellular egg) were distinguished in egg formation process (Fig. 2, □-3, □-3,4, □-2,3, □-2 and □-2,3). The oocyte was surrounded by follicular cells (Fig. 2, □-2, yellow arrows) and support cells (Fig. 2, □-2, SC) along the axis of the ovariole, revealing that oogenesis of P. *Artemia* followed a polytrophic oocyte mode.

**Fig. 2.**
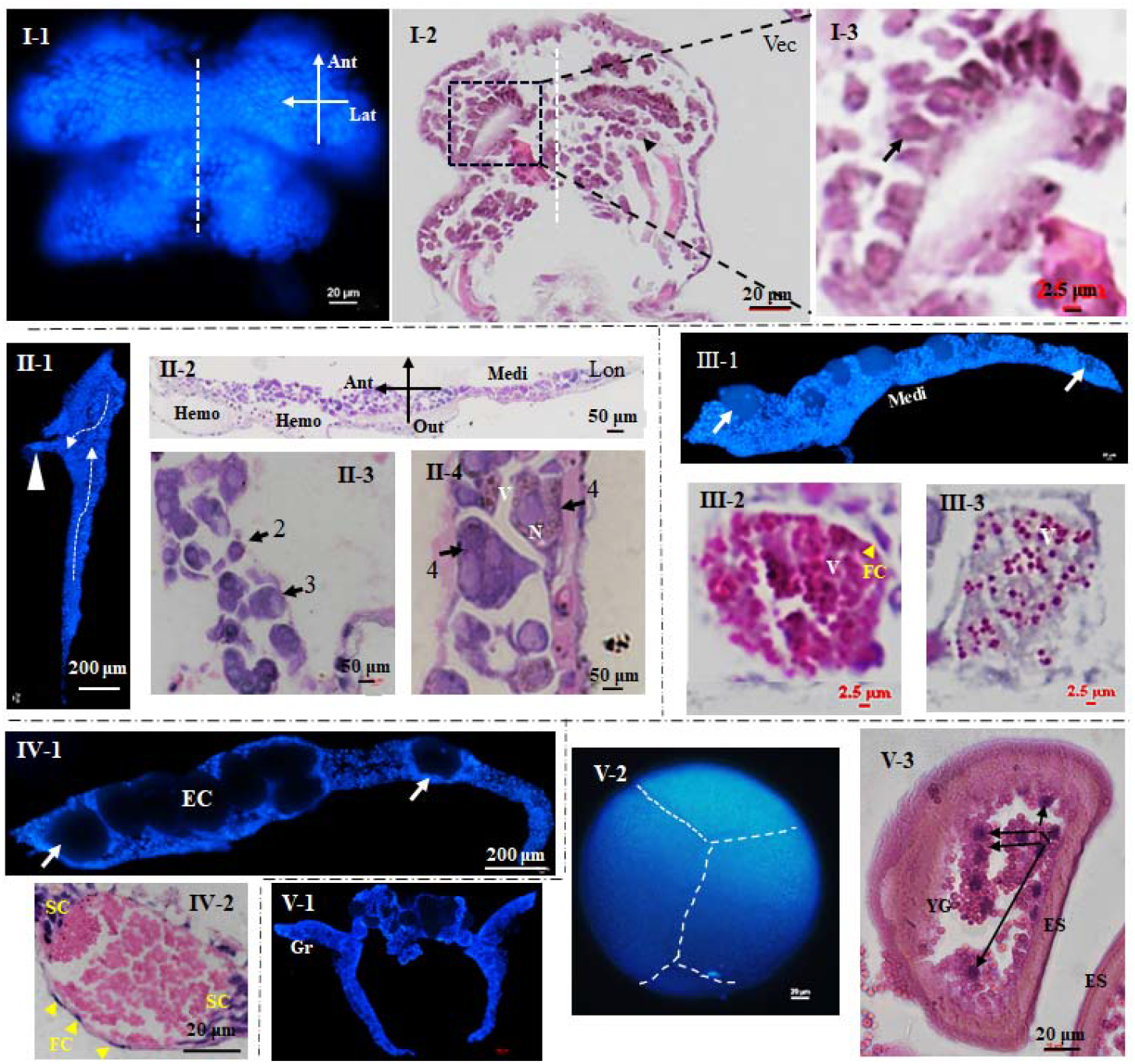
Cytological characteristics of oocytes accompanied by the ovarian development of P. *Artemia*. I-□ represented five stages of ovarian development. Stage □: The dark dots of square frame in Fig. □-2 is zoomed into Fig. □-3; dark arrows indicate the oogonia in ovariole; Vec: vector section. Stage □: white square dots in Fig. □-1 indicate the migration route of egg formation; Ant: anterior; Hemo: hemocoel; Lon: longitudinal section; Medi: medial surface; N: nucleus; Or: ovarioles; Out: outside. Stage □: FC: follicle cells (yellow arrows); V: vitellogenin. Stage □: SC: support cells; EC: egg chamber; Gr: germarium; white arrows indicate the oocytes in different regions of ovariole. Stage □: ES: egg shell; YG: yolk granules; white arrowhead in Fig □-1: oviduct channel.

At stage □, the ovary was not fully developed which was transparent with a small volume and lack of paired intact ovarioles (Fig. 2, □). The 1^st^ phase oocytes (oogonia) had a high nucleo-cytoplasmic ratio, nearly round shape and highly homogeneous by HE staining (Fig. 2, □-2,3, dark arrow). The 2^nd^ phase oocytes (Fig. 2, □-3, dark arrow) occurred at stage □ with irregular elliptic shape and bigger size. The 3^rd^ phase oocytes (Fig. 2, □-3, dark arrow) were round or polygonal with prominent nucleoli, and the 4^th^ phase oocytes (Fig. 2, □-4, dark arrow) had irregular shape with an increased cell volume. Significantly, the oocyte maturation process was characterized with gradually increased oocyte size and yolk granules, but decreased nucleo-cytoplasmic ratio (Fig. 2, □ to □).

At stage □, the 5^th^ phase oocytes were observed. As the ovary developed, the number of opaque eggs increased. The egg chamber (EC) gradually formed in the ovarioles and contained the eggs that about to maturity. Notably, the follicular cells (Fig. 2, Ш-2, FC, yellow arrow) were observed in stage IV, a row of single follicular cells (Fig. 2, □-2) surrounding to oocytes (Fig. 2C). At stage □, the 6^th^ phase oocytes were fully matured and entered the oocysts, which were enlarged black or gray cells. Before releasing by female, the cytoplasm cleavage occurred to form the multicellular egg (syncytium) and eventually develop into the gastrula embryo.

### Spatial dynamic changes of germline stem cells (GSCs)

As shown by immunofluorescence images, the ovary of P. *Artemia* included four parts (Fig. 1, Fig. 3): a pair of ovarioles (Or), oviduct (Ovd), oocyst (Oc) and ovulatory hole (Oh). The fat body (FB) was tightly embedded in the ovary (Fig. 3, B, C). By EdU labelling, the GSCs were identified (Fig. 3, A, B) during stage □-Ш of ovarian development. Oocytes at other different phases were confirmed according to the above results (Fig. 2) and their spatial position during ovarian development (Fig. 3). In stage □ and □ of ovarian development, the GSCs of P. *Artemia* were generated in the germarium (Gr), which contains dividing primordial germ cells and oogonia produced from them. Different from the generation, maturation and migration routes of *Drosophila* GSCs, the Gr of P. *Artemia* located at the proximal end of the oviduct (Fig. 3A, white dotted circles). The vitellarium (Vr) of P. *Artemia* located at the distal end of the oviduct (Fig. 3C). As a pair of ovarioles fully developed, the oocytes increased in number and were surrounded by follicular cells, which moved downward and migrated to the Vr where they continued to develop and mature (Fig. 3A). At this time, the Vr expands to form a series of egg chamber, in which oocytes gradually deposited into yolk protein. To complete the reproduction, the eggs (gastrula embryos) either develop into nauplii via ovoviviparous reproduction, or get surrounded by a thick shell (ES, Fig. 2V-3) and entered a state of diapause (cysts, oviparous reproduction), which are released through ovulatory hole (Fig. 1 triangular white arrow).

**Fig. 3.**
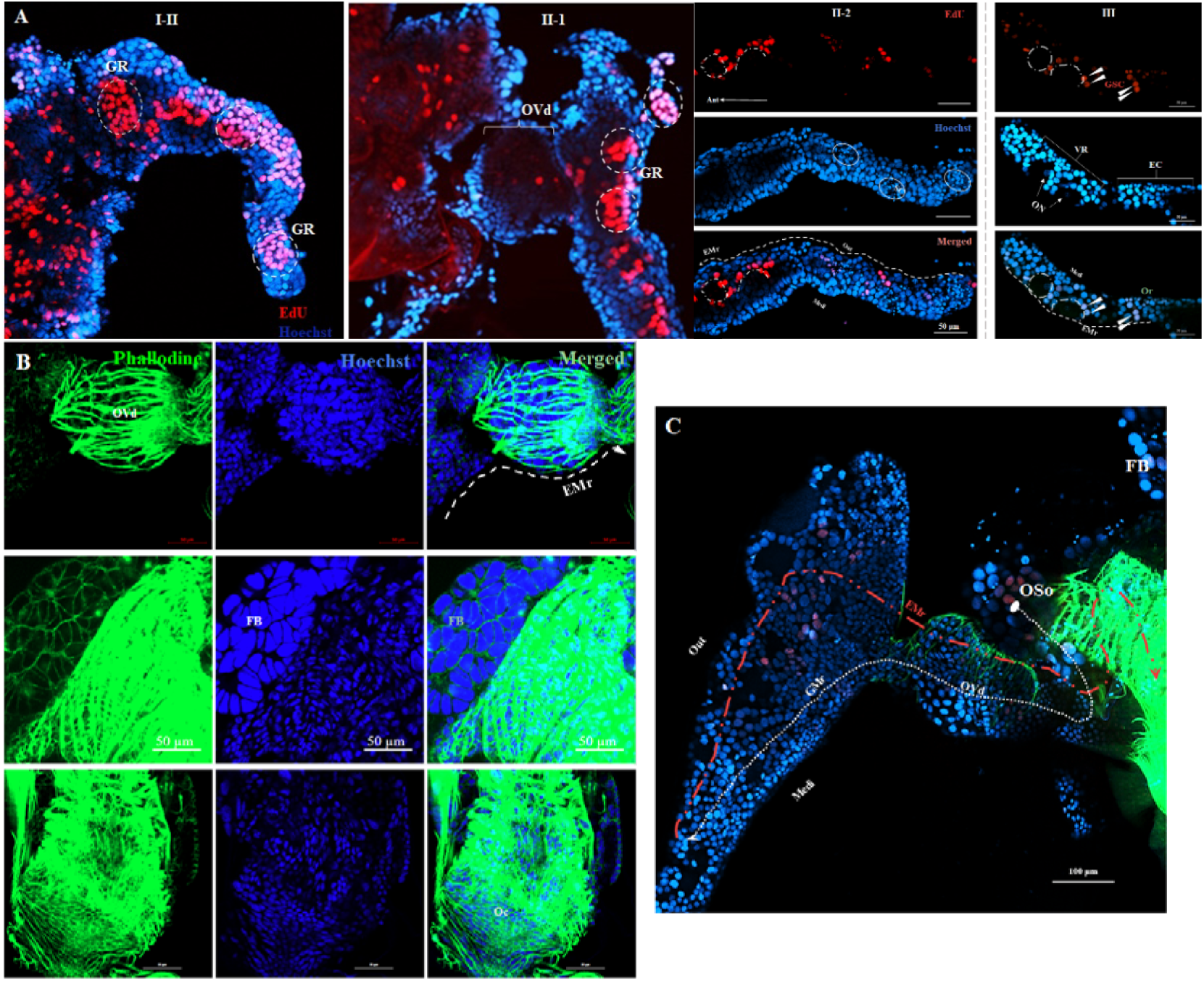
Spatial dynamic of germline stem cells in ovarian development. (A) GSCs translocation. White dot line indicates the direction; EC: egg chamber; EMr: egg formation and migration route; Gr: germarium; Vr: vitellarium; GSC: germline stem cell; white arrows indicate the GSCs; ON: oocyte nucleus; OP: oocyst primordium; SC: support cells; OI: oocyte I; Or: ovarioles. (B) Structural features of the ovary. FB: Fat body; Ovd: oviduct; Oc: oocyst. (C) Migration pathways of reproductive stem cells. GMr: germline stem cell migration route; white dot line indicates the direction; Medi: medial; Out: outward; Oso: original site of oogonia.

### Transcriptome analysis of ovaries at five developmental stages

In total 131,442 transcripts and 60,801 unigenes were obtained from the ovary tissues at five development stages (Table S1). The length of transcripts ranged from 301 bp to 19,780 bp, with an average length of 1,199 bp. And the length of unigenes ranged from 301 bp to 19,780 bp, with an average length of 1,077 bp. All sequences were spliced by Blast and compared with seven databases including Pfam, Nr, Swissprot, etc. to obtain corresponding annotation information. The results revealed that 14,816 unigenes were annotated by Nr library, 2.8% of which were similar to the matched sequences. Among unigenes matched with Nr database, 12.4% of them were *Daphnia magna* genome sequence, followed by *Daphnia pulex* (9.7%), termites *Crytotermes secundus* (4.6%), pea aphids *Acyrthosiphon pisum* (3.6%) and termites *Zootermopsis nevadensis* (3.3%) (Fig. 4). The annotations by other databases were shown in Table S2 and Fig. S1-S3.

**Fig. 4.**
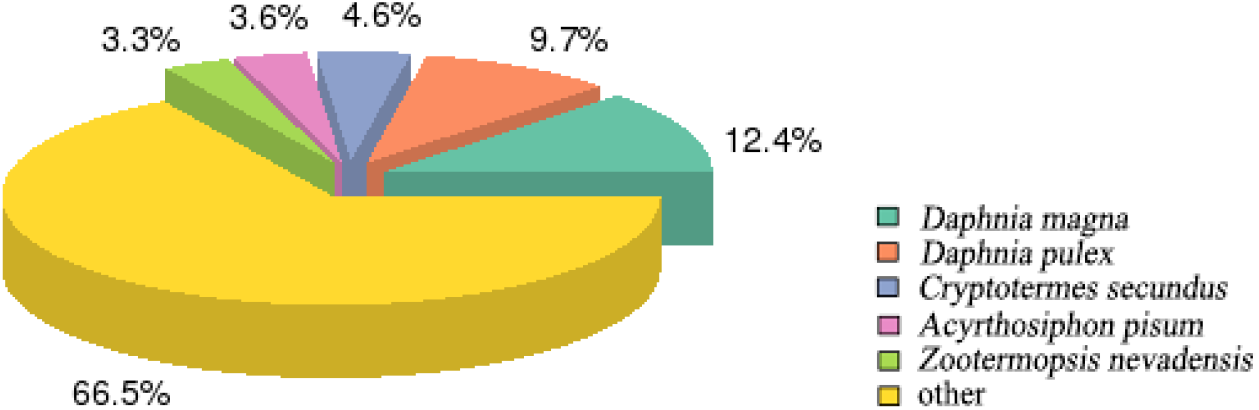
Statistical map of species distribution by Nr blasting.

### Differential expression gene analysis and KEGG analysis

To clarify the profiles of gene expression in ovary development process, 28459 differential expressed genes (DEGs) were identified by transcriptome at five ovarian developmental stages (Table 3 and Table 4). The number of DEGs in stage □ vs stage □ was 2,074, including 1,169 up-regulated genes and 905 down-regulated genes, while 1,116 DEGs were obtained in stage □ vs stage □, including 371 up-regulated genes and 745 down-regulated genes. Similarly, 948 DEGs were obtained in stage □ vs stage □, including 661 up-regulated genes and 287 down-regulated genes. The number of DEGs in stage □ vs stage □ was 819, including 456 up-regulated genes and 363 down-regulated genes.

**Table 3.**
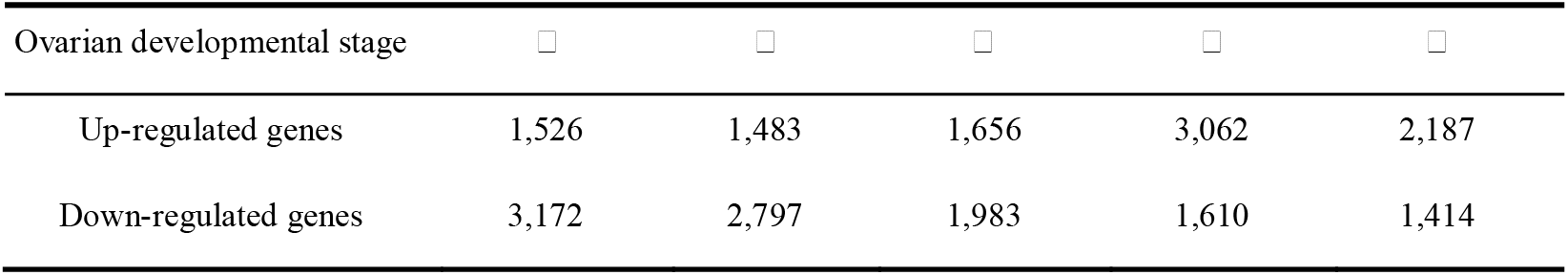
Variation in gene expression during the ovarian development of P. *Artemia*.

| Ovarian developmental stage | □ | □ | □ | □ | □ |
| --- | --- | --- | --- | --- | --- |
| Up-regulated genes | 1,526 | 1,483 | 1,656 | 3,062 | 2,187 |
| Down-regulated genes | 3,172 | 2,797 | 1,983 | 1,610 | 1,414 |

**Table 4.** Differential expressed genes during the ovarian development of P. *Artemia*.

| Ovarian developmental stage | Total | Up-regulated genes | Down-regulated genes |
| --- | --- | --- | --- |
| □ vs □ | 2,074 | 1,169 | 905 |
| □ vs □ | 1,116 | 371 | 745 |
| □ vs □ | 948 | 661 | 287 |
| □ vs □ | 819 | 466 | 363 |

To explore the function of key genes in the process of oogenesis, the DEGs obtained in ovarian development stage □ vs stage □ were statistically analyzed. Volcano map showed the number of common and unique DEGs in the two groups (Fig. 5A). KEGG pathway analysis showed that 121 pathways were enriched in stage □ vs stage □, and each pathway represented one or more biological processes. These pathways were relevant to progesterone-mediated oocyte maturation, cell cycle and oocyte meiosis which will provide important value for subsequent studies on reproductive development (Fig. 5B). The DEGs analysis in other ovarian developmental stages were shown in Fig. S4 and Fig. S5.

**Fig. 5.**
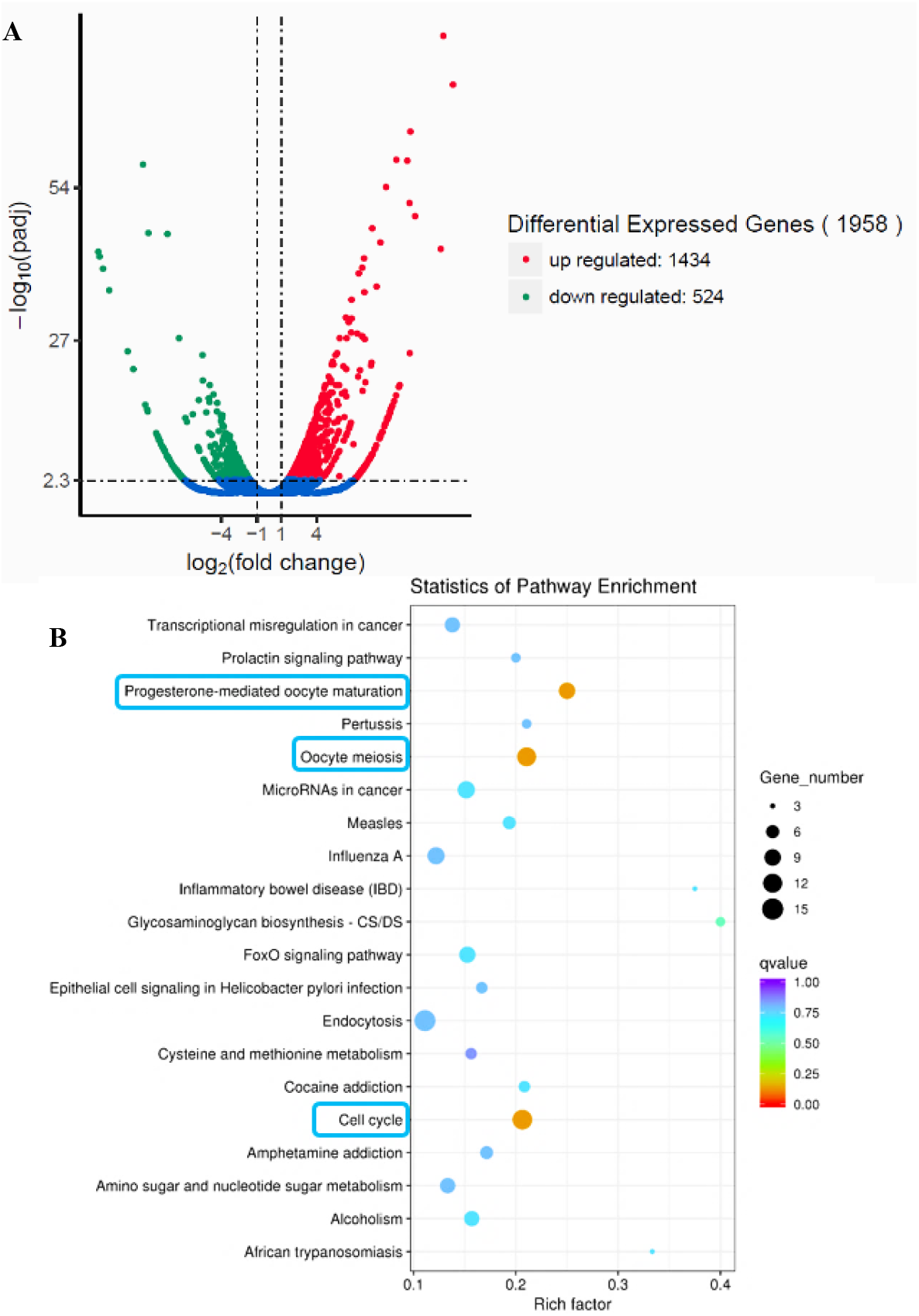
Volcanic map and enrichment pathway map of the two stages. Volcanic plot (A) and scattered plot of KEGG enrichment (B) of DEGs in the ovaries of P. *Artemia* in developmental stages IV vs stage II.

### Expression profile of transcripts involved in the reproduction-related genes

The FPKM value of each comparison group in the differential gene set of all comparison groups was calculated and were used for hierarchical clustering analysis of the DEGs expression level (Fig. 6A). Some reproductive DEGs were screened, and the key genes in BMP signaling pathway (*Dpp* and *Gbb)* and JH signaling pathway (*VgR* and *VgA)* and other reproduction-related genes (*vasa, piwi* and *nanos*) were further verified by qRT-PCR (Fig. 6B). It showed that the expression profiles of these genes were consistence with transcriptome sequencing results (Fig. 6C). At all sampling time, transcript expression levels varied among the seven genes. But higher gene expression level of *VgR, VgA, Dpp* and *Gbb* were obtained, peaking at stage □ and/or stage □. The expression profile of the above genes provided clues and basis f**B**or implementing of P. *Artemia* RNAi manipulation at the appropriate chance.

**Fig. 6.**
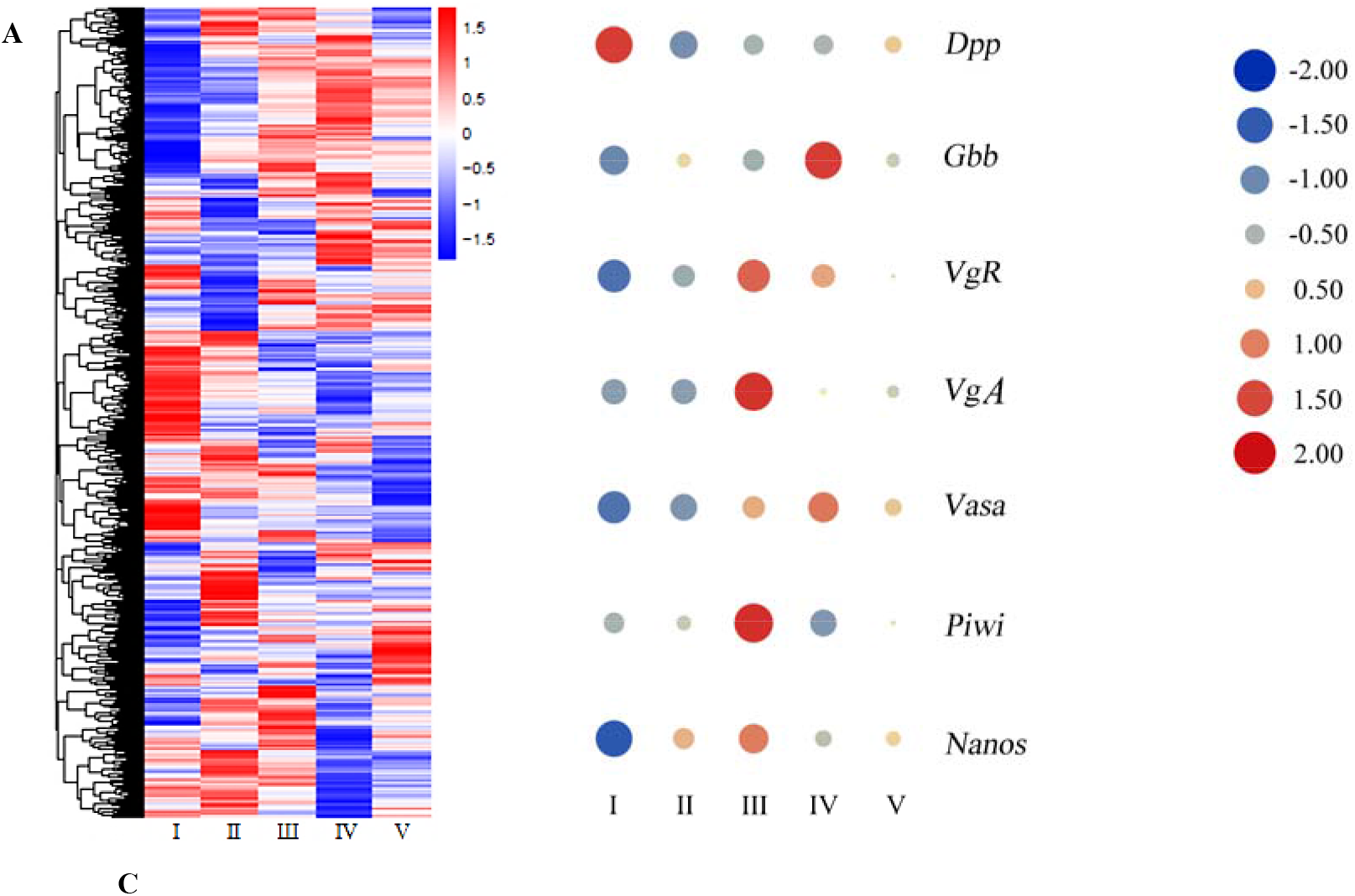
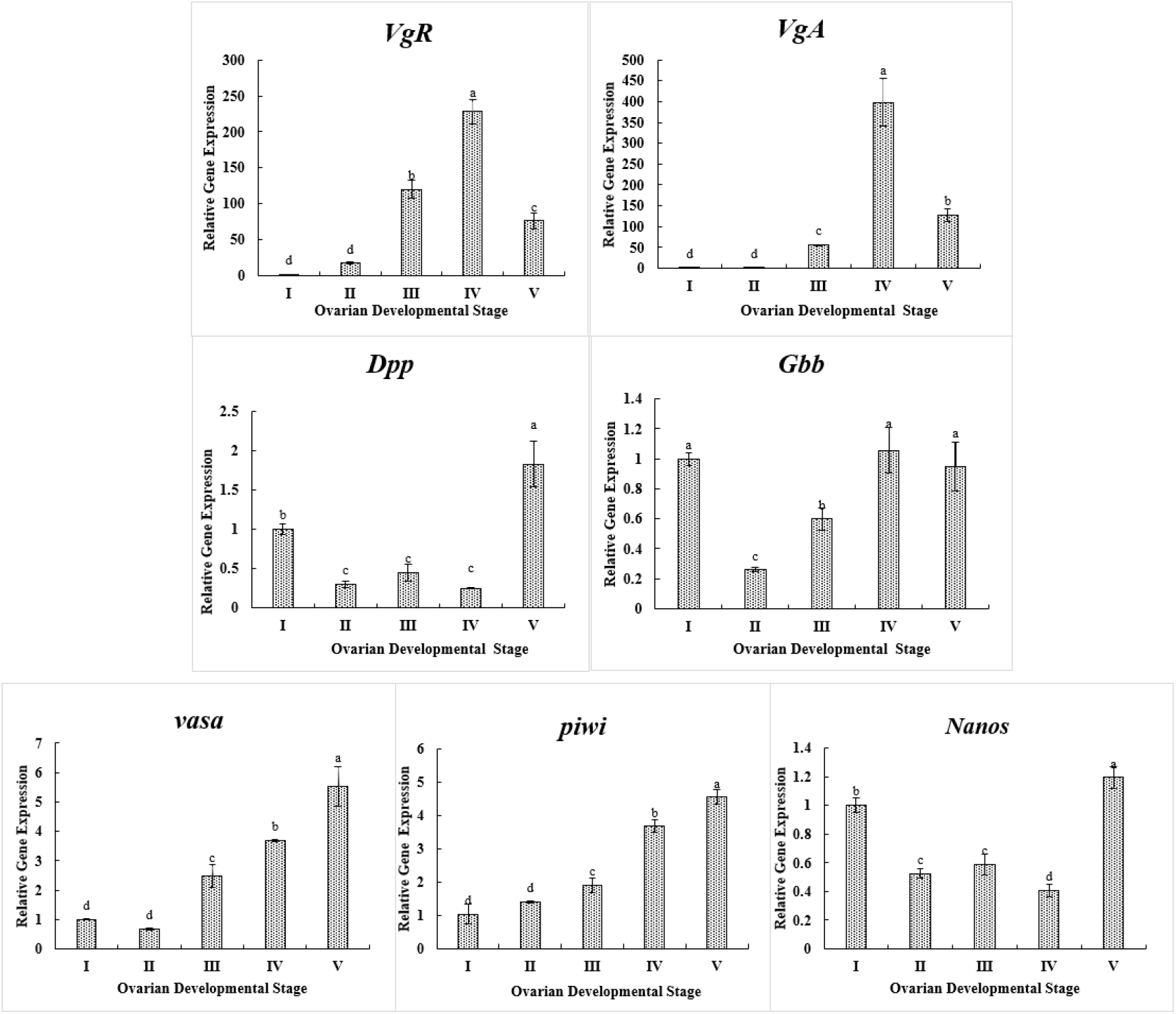
Analysis and qPCR verification of DEGs. (A) Clustering heat map of DEGs in ovaries at five developmental stages of P. *Artemia*. (B) Differential expression of the selected reproductive genes. (C) Variation of genes expression levels at five ovarian developmental stages of P. *Artemia*. Data are expressed as the mean ± SD (n = 30). Different alphabets on the bars represent the significant difference among the groups (P<0.05 or P<0.01).

### Structural characterization and phylogenetic analysis of *VgR* in P. *Artemia* (*PA-VgR*)

Similar to the *VgR*s of other crustaceans (Sun et al., 2020), *PA-VgR* contained three typically conserved modular elements for the Low-Density Lipoprotein Receptor (LDLR) superfamily (Fig. 7A). Ligand-binding domain (LBD) mediates receptor binding to ligands and contains class A repeats (LDLa). EGF-precursor homology domain (EGFD) mediated dissociation of receptor and liganded and consists of YWXD motifs and epidermal growth factor (EGF)-like repeats. At the carboxyl terminus, transmembrane domain (TM) formed a transmembrane α-helix, which anchored the receptor to the plasma membrane. But *PA-VgR* did not contain O-liked sugar domain (OLSD).

**Fig. 7.**
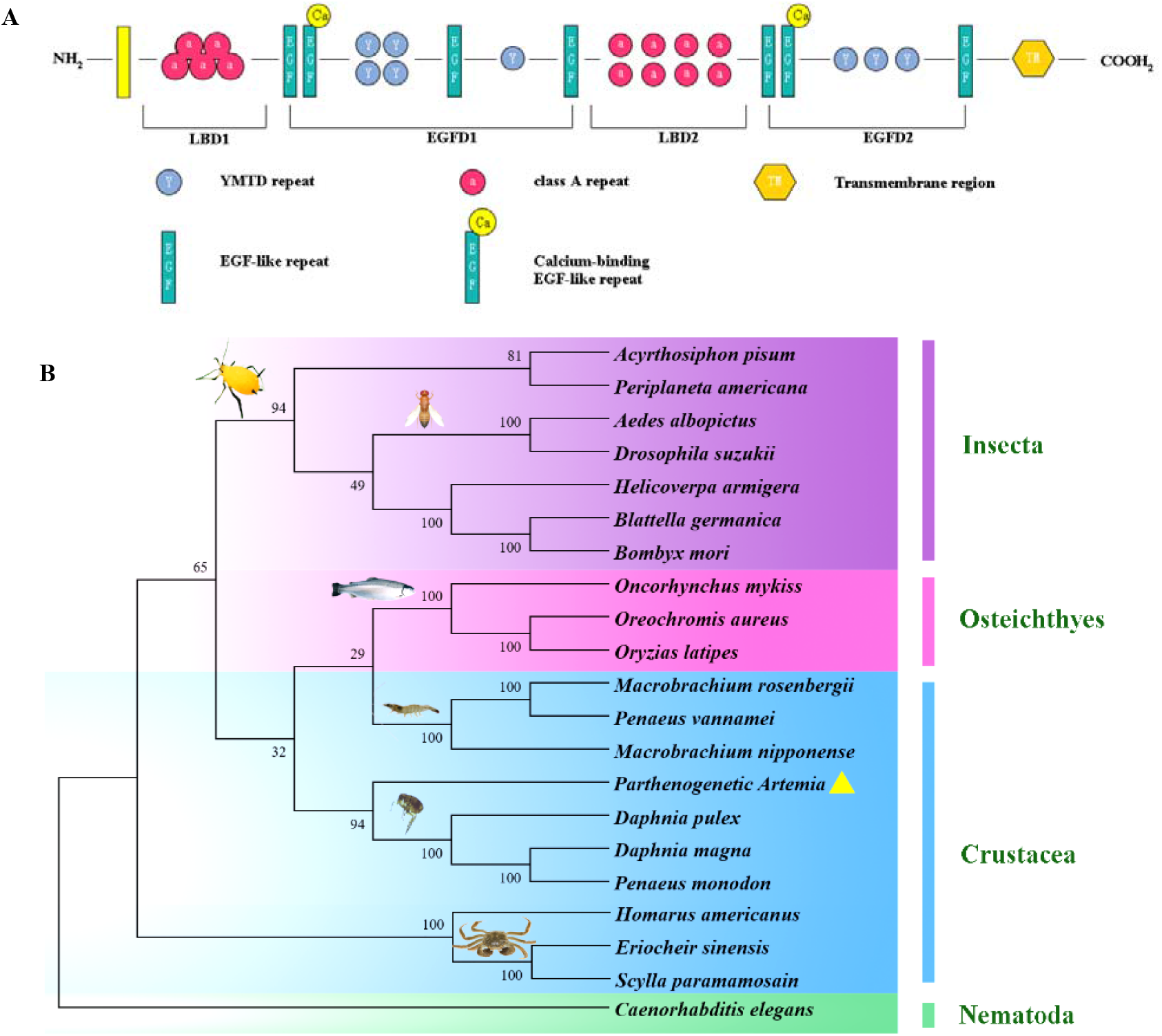
Protein structure of VgR in P. *Artemia* and phylogenetic analysis of *VgR*s in related species. (A) The structure domains of *PA-VgR* protein of P. *Artemia*, including ligand-binding domain (LBD), EGF-precursor homology domain (EGFD) and transmembrane domain (TM). (B) Phylogenetic tree of *VgR*s in various oviparous species.

The Genbank accession numbers of protein were shown in Table S3. The phylogenetic analysis showed that the *VgR*s from various oviparous species were grouped into three clades affiliating to vertebrates, arthropods and nematodes, and *PA-VgR* fitted in a clade of crustaceans and was more closely related to Daphniidae.

### Expression profile of corresponding genes after *PA*-*VgR* slicing

The transcript expression of *PA-VgR* were detected in the intestine, cerebral ganglion and ovary of P. *Artemia* by qPCR (Fig. 8A). The results showed that *PA-VgR* was specifically expressed in the ovary, but hardly detected in other two tissues. And the expression level of *PA-VgR* increased continuously and reached the highest at ovarian developmental stage IV. These results were consistent with the trends of RNAseq.

**Fig. 8.**
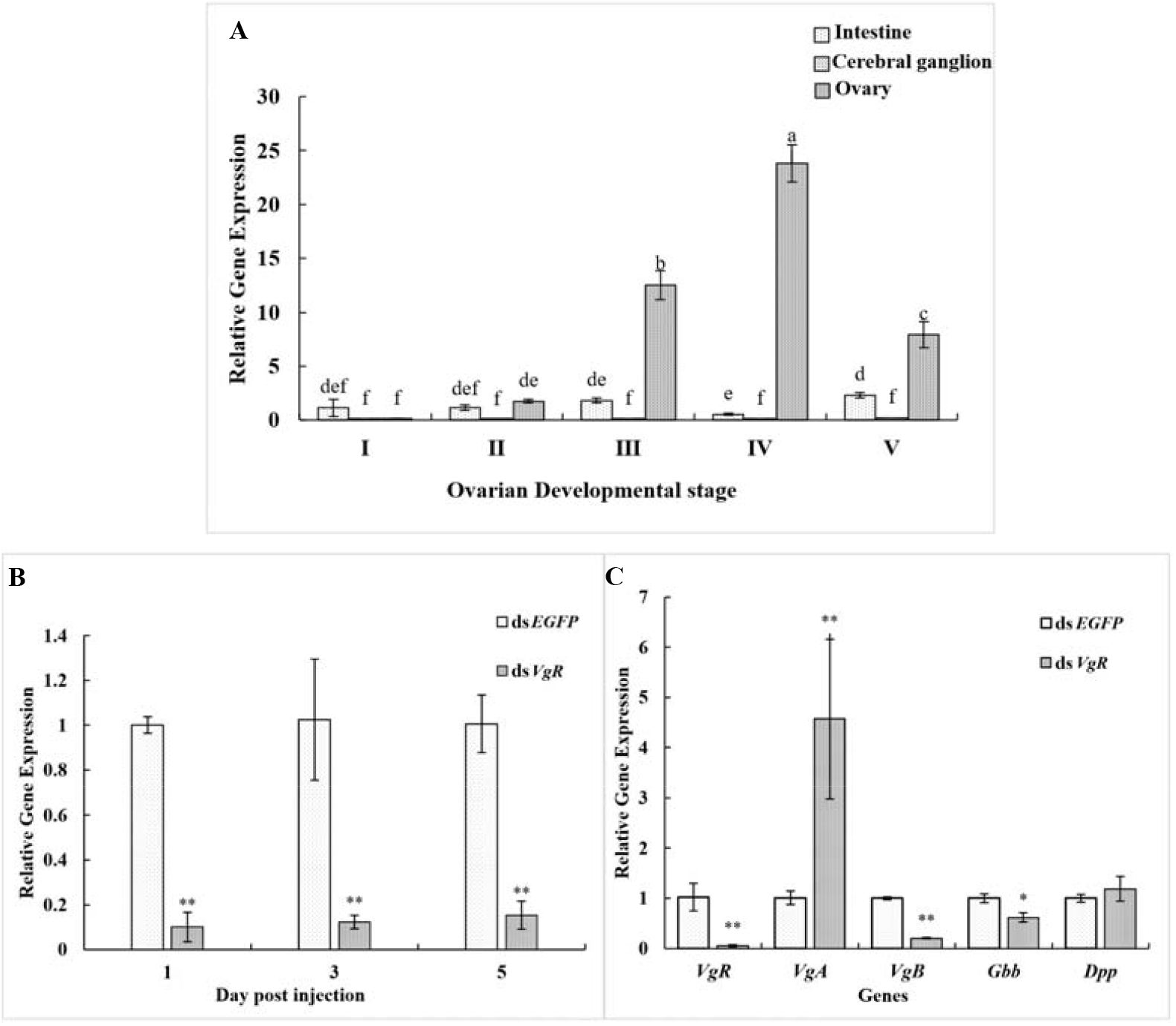
Profiles of *VgR* expression in process of ovarian development of P. *Artemia*. (A) Expression levels of *PA-VgR* mRNA in intestine, cerebral ganglion and ovary of at five ovarian development stages (I-V). (B) Expression level of *PA-VgR* mRNA on the 1^st^, 3^rd^, and 5^th^ day post injection of ds*EGFP* and ds*VgR*. (C) Expression levels of *VgA, VgB, Gbb* and *Dpp* mRNA on the 1^st^ day post dsRNA injection. Data are expressed as mean ± SD (n = 30). Different letters represent significant difference among the groups (P < 0.01). The “_*_” and “_**_”indicates significant difference between two groups (P<0.05, P<0.01).

In this study, RNAi assays were conducted to reveal the functions of *PA-VgR* in oogenesis of P. *Artemia*, and the dsRNA injection was performed at pro-mature stage according to transcriptome results (Fig. 5). Compared to ds*EGFP* group (control), the level of *PA-VgR* in ds*VgR*-treated group decreased by 89.8%, 90.2% and 85.2% on the 1^st^, 3^rd^, 5^th^ day post injection, respectively (Fig. 8B). The effect of *PA-VgR* RNAi on the expression of oogenesis-related genes *VgA, VgB, Dpp* and *Gbb* were further studied. Compared to ds*EGFP* group, the expression of *VgA* was significantly increased by silencing *PA-VgR*. The expression of *VgB* and *Gbb* was down-regulated, while the expression of *Dpp* was not significantly changed (Fig. 8C).

### Phenotypes modulation of oogenesis after *PA-VgR* knockdown

The ovaries of *PA-VgR* knockdown individuals exhibited clear phenotypic changes (Fig. 9A). *In vivo* observation revealed that the egg development in ovary was delayed on the 3^rd^ day after ds*VgR* injection. On the 5^th^ day after ds*VgR* injection, the abnormal egg development, disordered distribution of eggs in ovarioles and reduced number of eggs were observed. *In vitro* observation showed that the ovaries of ds*VgR* group had a large number of deformities and less opaque (Fig. 9B, lower right figure). The biometric measurement of the ovarioles showed that the P. *Artemia* in ds*VgR* group had significantly shorter length of the ovarioles, and shorter width of the germarium, oviduct and vitellarium (P<0.05), which further proved that the knockdown of *PA-VgR* gene delayed the ovarian development and ultimately led to the reduced fecundity (Fig. 9B, C and Table 5).

**Fig. 9.**
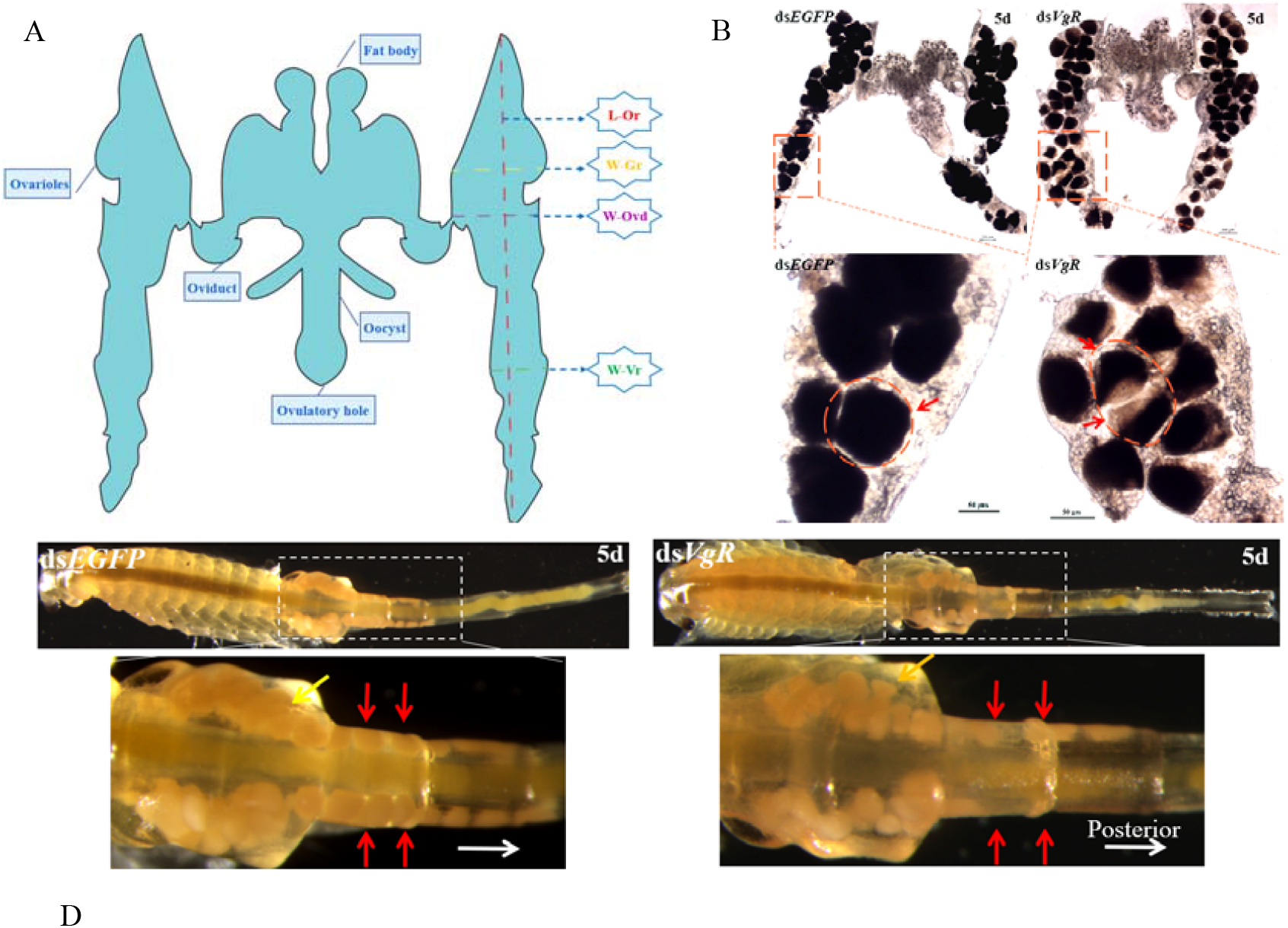
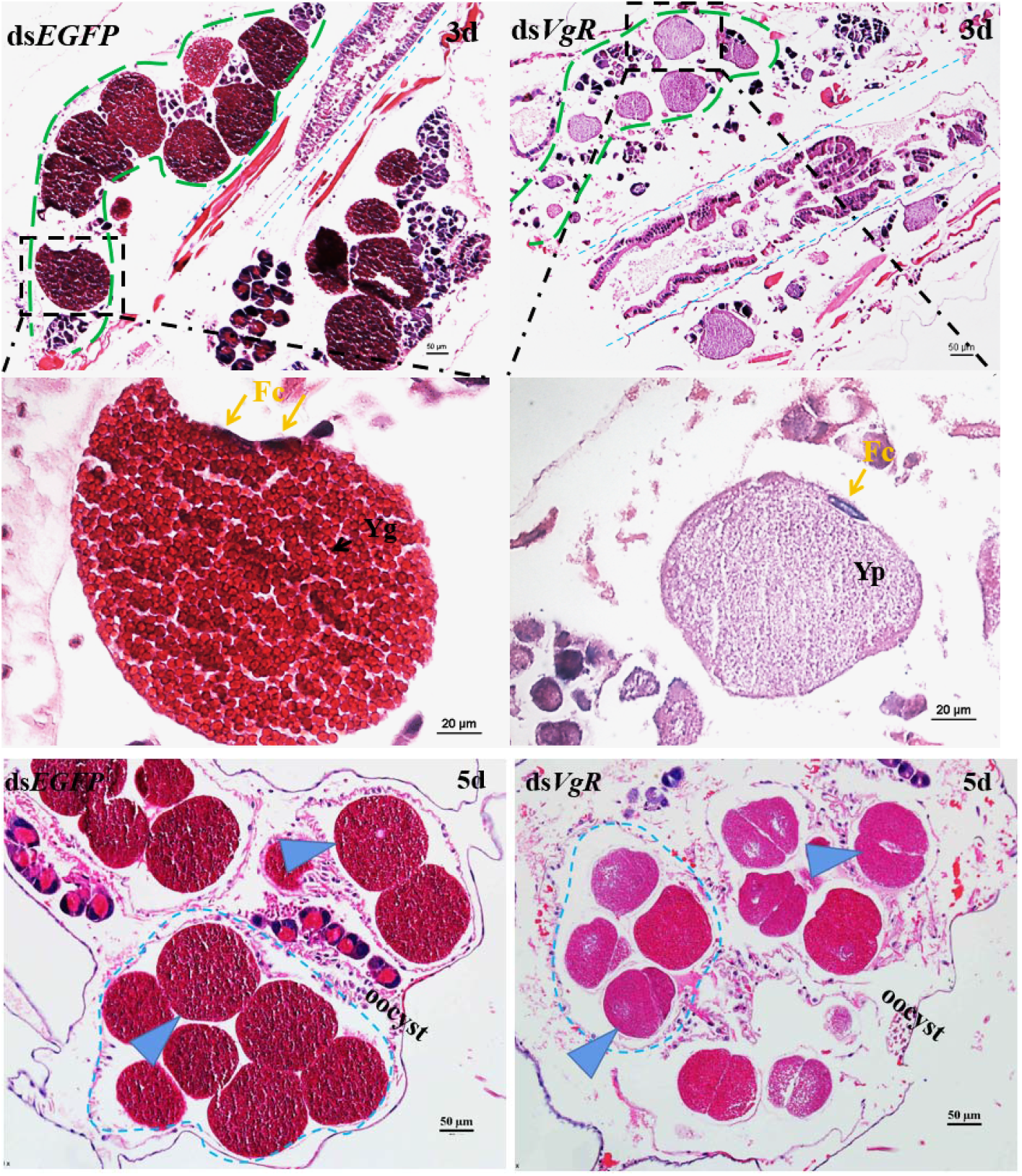
Anatomical and morphological feature of P. *Artemia* ovaries after *PA-VgR* silencing. (A) Schematic figure of mature P. *Artemia* ovary. L-Or: Length of the ovarioles; W-Gr: Width of the germarium; W-Ovd: Width of the oviduct; W-Vr: Width of the vitellarium. (B) *In vitro* observation of P. *Artemia* ovary in ds*VgR* and ds*EGFP* group. Red arrows indicate the eggs. (C) *In vivo* observation of P. *Artemia* ovary in ds*VgR* and ds*EGFP* group. The white arrows dots showed migration route of egg formation. Red arrows indicate the eggs distributing symmetrically in ovarioles. (D) Cross section of HE stained ovaries. Green arrows indicate the eggs in ovariole. Yellow arrows indicate the follicular cells.

**Table 5.** Biometric measurement of P. *Artemia* ovarioles after RNAi (n=10)

| | Length of the ovarioles<br>($\mu\text{m}$) | Width of the germarium<br>($\mu\text{m}$) | Width of the oviduct<br>($\mu\text{m}$) | Width of the vitellarium<br>($\mu\text{m}$) |
| --- | --- | --- | --- | --- |
| *dsEGFP* group | 1484.46±226.71 | 232.93±70.39 | 233.63±44.35 | 193.89±92.64 |
| *dsVgR* group | 1384.07±223.71 | 214.94±119.24 | 149.19±45.73* | 134.42±75.34 |

On the 3^rd^ day post dsRNA injection, HE staining of ovary showed no obvious morphologic changes in follicular cells after *PA-VgR* gene silencing, and all of them had formed unicellular eggs (Fig. 9D). Meanwhile the yolk protein was further accumulated and aggregated in the eggs to form a large amount of yolk granules in the ds*EGFP* group (Fig. 9D), whilst the eggs in ds*VgR* group were deformed with less accumulation of yolk granules but rich in yolk protein. On the 5^th^ day after dsRNA injection, the accumulation of yolk granules in ds*EGFP* group was completed and the typical granular eggs were formed in the oocyst. While knocking down *VgR* gene, cleavage achieved in advance with obviously uneven and a large number of vacuoles, also indicated the inferior egg quality. Meanwhile the number of eggs in ds*VgR* group was significantly lower than that in ds*EGFP* group (P<0.01) on both 3^rd^ and 5^th^ day after ovulation period (Fig. 10 and Table 5).

**Fig. 10.**
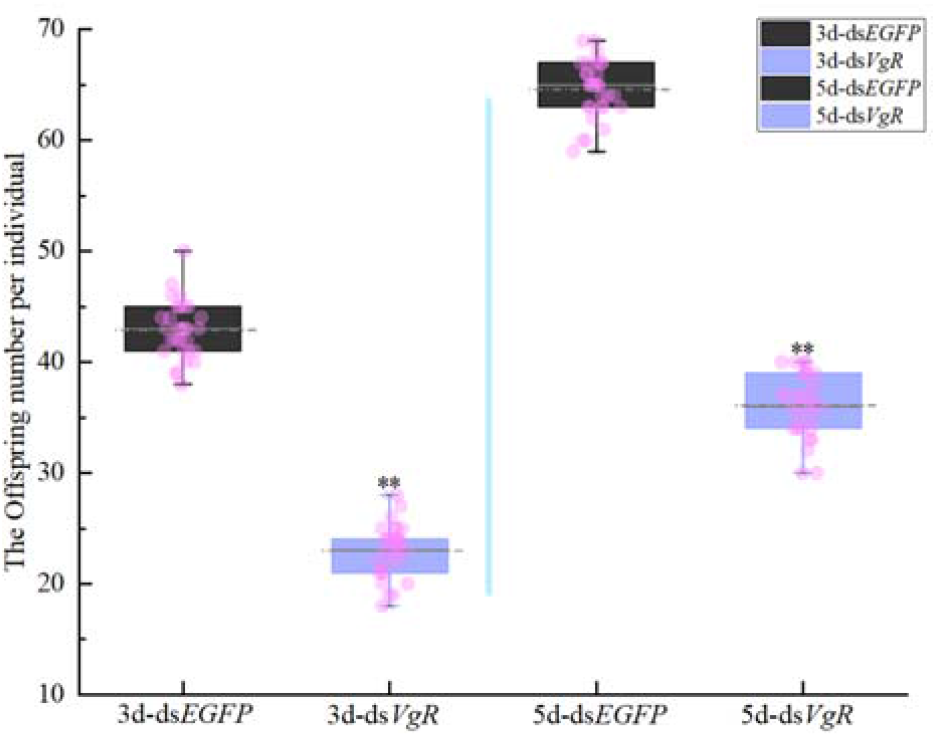
Offspring number in per individual of P. *Artemia* on the 3^rd^ and 5^th^ days after the ovulation period (n=30). The double asterisk (**) indicates a significant difference between two groups (P<0.01).

## DISCUSSION

The continuation of life from generation to generation through reproduction and development is the biological base of life proliferation (Johnson et al., 2001). The biological advantages of P. *Artemia*, such as colonial offspring production, relatively short lifespan and reproductive cycle, enable it more suitable for the reproduction process and molecular mechanism study (Gay et al., 2018; Wang et al., 2021). In this study, the spatio-temporal features of ovarian development and oogenesis in P. *Artemia* were clarified by means of *in vivo* and *in vitro* approaches. Notably, fat body tissue was firstly identified in *Artemia*, which is assumed to play a coincident role in promoting vitellogenesis as fat body of insects and the hepatopancreas of crustaceans (Souza et al., 2017; Wu et al., 2018). As known, both fat body and hepatopancreas provide vitellogenin and promote its binding for vitellogenin receptors. Moreover, we found some significant differences in generative sites of GSCs and migration paths between P. *Artemia* and *Drosophila* (Fu et al., 2015a; Shapiro-Kulnane et al., 2015). Upon the ovarioles development complete, GSCs of *Drosophila* generate from the germarium near to terminal filament (Donoughe et al., 2014), whilst the oogonia of P. *Artemia* originated from a pair of ovariole buds. Along with ovarian development, the GSCs migrated in the form of cell mass to the corresponding sites where niches regulate the egg formation (Fig. 1 and Fig. 3A). The formation of cell mass and GSCs niche in P. *Artemia* may be facilitated to the successive brood offspring production of *Artemia*, in favor of population sustainability in extreme environments.

The hormone- and gene-mediated regulation of reproductive development have been well studied in insects (Song et al., 2018; Guo et al., 2019), but the mechanism of reproductive development of crustacean in extreme environments are unexplored. The transcriptome platform is an important and convenient tool to provide data supporting the gene and gene network regulation (Carmona et al., 2017; Stavely et al., 2023). Our data revealed that 28,459 DEGs relating to ovarian development and oogenesis were explored in P. *Artemia*, in which the genes relevant to cell cycle and oocyte meiosis, the signaling pathway of insulin and HIF were significantly expressed, indicating that these may regulate and participate some key cyto-events in ovariole formation, GSCs generation and migration, vitellogenesis and choriogenesis (Armstrong and Drummond-Barbosa, 2018). In this study, *PA-VgR* were screened and verified according to the DEGs of the transcriptome. *VgR* has been confirmed a key receptor for vitellogenesis, which is fundamental for oocyte development in oviparous animals and serves as a key preparatory stage for establishing energy reserves and biomolecules for subsequent embryonic development (Jorgensen et al., 2009; Shang et al., 2018).

We obtained a full-length cDNA of *VgR* in P. *Artemia* and predicted its corresponding protein structural domains (Fig. 7A). Similar to other members in the LDLR superfamily, *PA-VgR* contained three highly conserved regions (LBD, EGFD, TM). However, similar to some insects such as fire ants (Chen et al., 2004) and *Drosophila melanogaster* (Schonbaum et al., 1995), there was lack of O-linked Carbohydrate Domain (OLSD) on the plasma membrane surface in P. *Artemia* VgR. It has been demonstrated that the tissue expression specificity of *VgR* is related to the existence of OLSD structure in VgR (Pousis et al., 2012). In this study, the expression level of *PA-VgR* without OLSD was significantly higher in ovary than in other tissues, while the expression level of *VgR* in other species with OLSD was higher in non-ovarian tissues (Lee et al., 2014; Lu et al., 2015). It has also been found that natural absence of VgR in OLSD can cause familial hypercholesterolemia in humans (Koivisto et al., 1993). Therefore, OLSD structure in VgR can be a potential marker indicating reproductive specificity of species.

It has been reported that RNAi of *VgR* suppressed Vg accumulation in the ovary of tiger shrimp *P. monodon* (Tiu et al., 2008b) and delayed maturation of the ovary in freshwater shrimp *M. nipponense* (Bai et al., 2016). However, the phenotypic details on *VgR* regulation in process of ovarian development were not well understood and controversial in different crustacean species (Tiu et al., 2008a; Bai et al., 2016). In our study, the expression of *PA-VgR* was specifically detected in the ovary of P. *Artemia* (Fig. 8), which is consistent with most insects (Liu et al., 2011; Smith and Kaufman, 2013). Our results illustrated that the expression levels of *PA-VgR* in the ovary increased continuously from the ovarian developmental stage I to stage IV (Fig. 4B), indicating that a exogenous Vg absorption process similar to other oviparous animals which exhibit exogenous vitellogenesis, such as fish (Hiramatsu et al., 2015) and insects (Zhu et al., 2020). However, the expression levels of *PA-VgR* reduced rapidly at ovulatory period (stage □), which was in accordance with increased yolk consumption. In fact, these yolk reserve will further sustain the survive of non-feeding *Artemia* nauplii for 3-4 days post hatching.

In this study, the possible molecular mechanisms underlying the onset of vitellogenesis and offspring production was firstly demonstrated by *PA-VgR* regulation in P. *Artemia*. The abnormal eggs resulted by ds*VgR* injection and relatively low *PA-VgR* expression for a period of time indicated that *PA-VgR* knockdown had a sustained inhibitory effect on yolk granules formation and eventually led to a significantly decreased offspring production (Fig. 10). Moreover, multiple copies of Vgs genes were screened from the transcriptome and named with capital letters. Our results indicated *PA-VgR* knockdown significantly increased the *VgA* expression and decreased *VgB* expression. As it has been reported in insects, *VgR* silencing might lead to the blocking of the expression of its ligand *VgB* (Jing et al., 2021), while the expression of *VgA* gene greatly increased, which seemed to be a compensatory of Vg synthesis to restore its nutrient reserve as soon as possible (Tiu et al., 2008b). Another explanation is that different *Vg* gene family member involved in different physiological functions, like yolk protein formation and plays non-nutritional roles (Zhang et al., 2015). It is necessary to investigate the molecular characteristics of vitellogenin family as well as their functions in ovarian development, oogenesis, immune response or nutrient delivery. On the other hand, *PA-VgR* silencing reduced the expression of *Gbb*, a key gene in BMP signaling pathway. BMP plays an regulatory role in the embryonic development of mice (Yu et al., 2010) and the proliferation, differentiation and regeneration of germline stem cells in *Drosophila* (Wotton et al., 2013), and metamorphosis of insects (Ishimaru et al., 2016). We propose that the decrease of *Gbb* expression affected the development and offspring production by interfering the normal proliferation and differentiation of germline stem cells.

In conclusion, our study revealed the process and the regulation mechanism of ovarian development and the egg formation of P. *Artemia*. More interesting, the structure and function of *PA-VgR* gene were more similar to insects, but different from other crustaceans of the same Class. We speculate that P. *Artemia* had differentiated into the same class as crustaceans in ancient times, but some physiological characteristics and functions might be more similar to insects along with the evolution and the adaptation to the extreme environment. Furthermore, the assay of *VgR* gene silencing lay the foundation for *in vivo* gene editing in *Artemia*. With the publication of the genome of *Artemia franciscana* (Yuan et al.; De Vos et al., 2021) and the completion of the whole genome sequencing of P. *Artemia* (data not shown, ARARC sequenced), it is possible to find more accurate and comprehensive molecular evidence to interpretate the role of *Artemia* in the evolution of species at the omics level.

## Acknowledgements

We would like to thank Dr. Zengyu Ma for her help with qPCR and RNAi assays.

## Author contributions

Conceptualization: H. Duan, LY. Sui, JH. Xiang, XX. Shao, Y. Li; Methodology: H. Duan, XX. Shao, Y. Li, W. Liu; Software: XX. Shao, GR. Du; Validation: H. Duan, XX. Shao, XH. Wang; Formal analysis: H. Duan, XX. Shao, W. Liu, NM. Pan, JP. Zhou; Investigation: H. Duan, XX. Shao, JH. Xiang, Y. Li, W. Liu; Resources: XX. Shao, NM. Pan, LY. Sui; Data curation: H. Duan, XX. Shao, LY. Sui; Writing - original draft: XX. Shao, H. Duan; Writing - review & editing: H. Duan, XX. Shao, LY. Sui, JH. Xiang; Visualization: H. Duan, XX. Shao, XH. Wang, JP. Zhou; Supervision: H. Duan, LY. Sui, JH. Xiang; Project administration: H. Duan, XH. Wang; Funding acquisition: H. Duan.

